# Experimentally Evolved *Staphylococcus aureus* Survives in the Presence of *Pseudomonas aeruginosa* by Acquiring Mutations in the Amino Acid Transporter, GltT

**DOI:** 10.1101/2023.07.24.550428

**Authors:** Ashley M. Alexander, Justin M. Luu, Vishnu Raghuram, Giulia Bottacin, Simon van Vliet, Timothy D. Read, Joanna B. Goldberg

## Abstract

*Staphylococcus aureus* and *Pseudomonas aeruginosa* are the most common bacterial pathogens isolated from cystic fibrosis (CF) related lung infections. When both of these opportunistic pathogens are found in a coinfection, CF patients tend to have higher rates of pulmonary exacerbations and experience a more rapid decrease in lung function. When cultured together under standard laboratory conditions, it is often observed that *P. aeruginosa* effectively inhibits *S. aureus* growth. Previous work from our group revealed that *S. aureus* from CF infections have isolate-specific survival capabilities when cocultured with *P. aeruginosa*. In this study, we designed a serial transfer evolution experiment to identify mutations that allow *S. aureus* to adapt to the presence of *P. aeruginosa*. Using *S. aureus* USA300 JE2 as our ancestral strain, populations of *S. aureus* were repeatedly cocultured with fresh *P. aeruginosa* strain, PAO1. After 8 coculture periods, *S. aureus* populations that survived better in the presence of PAO1 were observed. We found two independent mutations in the highly conserved *S. aureus* aspartate transporter, *gltT*, that were unique to evolved *P. aeruginosa*-tolerant isolates. Subsequent phenotypic testing demonstrated that *gltT* mutants have reduced uptake of glutamate and outcompete wild-type *S. aureus* when glutamate is absent from chemically-defined media. These findings together demonstrate that the presence of *P. aeruginosa* exerts selective pressure on *S. aureus* to alter its uptake and metabolism of key amino acids when the two bacteria are cultured together.

**Importance:** *Staphylococcus aureus* and *Pseudomonas aeruginosa* are the two most common bacterial pathogens that infect people with the genetic disease, cystic fibrosis (CF). They are often found together in CF-associated polymicrobial infections that are associated with worse patient prognosis. Understanding how these very different opportunistic pathogens influence each other in a shared environment is pertinent to improving the treatment of polymicrobial infections. While much attention has been brought to the interspecific interactions between *S. aureus* and *P. aeruginosa,* few studies have used experimental evolution methods to identify determinants of their competition and coexistence. Here, we use a serial transfer experimental evolution approach and identified a single genetic change associated with improved survival of *S. aureus* in the presence of *P. aeruginosa.* Our findings implicate metabolism of shared resources as an important factor in *S. aureus’s* ability to survive in the presence of *P. aeruginosa*.

## Introduction

Often, a community of microbes contributes to the shaping of the infection environment through metabolic interactions, altering adaptive trajectories, or changing the antibiotic susceptibilities of interacting strains (Brown et al., 2008, 2012; Dalton et al., 2011; Stacy et al., 2016; Wadsworth et al., 2018). In the case of chronic infections, multiple opportunistic pathogens can coexist in their shared host environment for many generations, exerting their own respective selective pressures on each other (Hamelin et al., 2019). Pairwise interactions between coexisting opportunistic pathogens such as *Staphylococcus aureus* and *Pseudomonas aeruginosa* have become a topic of great interest to microbiologists both for the importance of the interaction to the course of the genetic disease cystic fibrosis (CF) and as model system for pathogen coevolution (Barraza & Whiteley, 2021; Camus et al., 2020).

In 2021, 32,100 people were documented as living with CF in the United States, a genetic disease that impacts multiple organ systems and greatly reduces life-expectancy and requires lifelong treatment (Cystic Fibrosis Foundation, 2021). One of the major complications of this disease is an increased risk for developing chronic respiratory infections that are exacerbated by the buildup of respiratory sputum. Over the last decade, *S. aureus* has displaced *P. aeruginosa* as the most common infective agent responsible for respiratory infections in people with CF and is detected in as many as 70% of CF associated lung infections. *S. aureus* is the predominant pathogen infecting young people with CF; older patients are more likely to be infected with *P. aeruginosa* and many individuals maintain both pathogens in coinfections (Cystic Fibrosis Foundation, 2021). Coinfections with *P. aeruginosa* and *S. aureus* may persist in the same patient for many years and even decades (Bernardy et al., 2020; Camus et al., 2020; Fischer et al., 2020). Chart reviews of more than 200 patients have revealed that coinfected patients experience significantly more pulmonary exacerbations and a more rapid decline in lung function compared to those with monoinfections of *S. aureus* or *P. aeruginosa* (Limoli et al., 2016).

When *P. aeruginosa* and *S. aureus* are cultured together outside of a host, there are a range of outcomes that may be dependent on strain identity or environmental conditions (Bernardy et al., 2022; Filkins et al., 2015; Limoli et al., 2017; Mashburn et al., 2005). Previously, our group observed that *S. aureus* isolated from CF patients range from highly sensitive to tolerant in their ability to coexist with the lab-adapted strain of *P. aeruginosa,* PAO1. Sensitive clinical isolates experienced as much as a 6-fold decrease in recovered colony-forming units (CFUs) after coculture, while others maintained most of their population, experiencing less than a 2-fold decrease in population size (Bernardy et al., 2020).

We have also observed that *S. aureus* is able to adapt to the presence of *P. aeruginosa* in its environment by showing that co-isolated *S. aureus* and *P. aeruginosa* strains grow better compared to non-concurrent isolates (Bernardy et al., 2022). Additionally, other groups have found that fermentative metabolism, polysaccharide production, and toxin excretion are all important phenotypes for *S. aureus* in its coexistence with *P. aeruginosa* (Filkins et al., 2015; Wieneke et al., 2021). It is also known that *S. aureus* can adapt to *P. aeruginosa* bactericidal compounds such as 2-heptyl-4-hydroxyquinoline-N-oxide (HQNO) and pyocyanin and that such adaptations may impact antibiotic resistance profiles of either species (Filkins et al., 2015; Limoli et al., 2016; Nguyen et al., 2014; Orazi & O’Toole, 2017).

In this study we sought to gain a greater understanding for how *S. aureus* adapts to the presence of *P. aeruginosa* and which *S. aureus* genotypes/phenotypes are under strong selective pressure in their shared environment. We showed that *S. aureus* adapted to the negative selective pressures presented by *P. aeruginosa* in a serial transfer evolution experiment. We found that rather than adapting to secreted toxins or contact dependent killing, *S. aureus* reduced its uptake of aspartate by disrupting its singular aspartate transporter, *gltT* (Potter et al., 2020). We hypothesize that loss of function of this membrane transporter results in *S. aureus* becoming more resilient to fluctuations in nutrient availability caused by the presence of *P. aeruginosa* in its environment. These results are surprising given that *P. aeruginosa* has other well-characterized mechanisms for directly inhibiting *S. aureus* in its environment, however, our findings suggest that optimizing amino acid metabolism is a potential pathway for adaptation for *S. aureus* that co-occurs with *P. aeruginosa*.

## Methods

### Bacterial Strains

*Staphylococcus aureus* USA 300 strain JE2 (Kennedy et al., 2008, 2010) was used as the ancestral strain for experimental evolution with *Pseudomonas aeruginosa* strain PAO1. Using the Nebraska Transposon Mutant Library (NTML) (Fey et al., 2013)an isogenic JE2 *gltT* mutant (*SAUSA300_2329)* was obtained and transduced into our own JE2 background. To do this we amplified the *gltT* locus with the transposon from *SAUSA300_2329*. The *gltT* gene with the inserted transposon was then confirmed and amplified by PCR and then transduced into our parental JE2 strain using phage φ11 (Iandolo et al., 2002) to generate strain JE2 *gltT*::Tn, as described below.

In brief, transduction was carried out by first preparing fresh φ11 lysate first with *S. aureus* strain RN4220. Collected lysate was then inoculated with NTML isolate *SAUSA300_2329* and titers were measured at 3×10^8^ pfu/mL. Transduction was then carried out with a multiplicity of infection of 0.1 according to methods in Krausz & Bose (2016). Transduced colonies were isolated on Trypticase Soy Agar (TSA) plates with 25µg/mL erythromycin and confirmed by PCR, identifying the presence of the complete transposon at the correct site on the chromosome. Primer sequences used to confirm transposon by amplicon size – 5’ AAAATTAGCCTACCTATGCAAGTTGT 3’ and 5’ TTTTGCTTTGTCATATACGTTTTCC 3’. We also used transposon specific primers to amplify from within the transposon and the gene itself using primers (negative strand) 5’ GCTTTTTCTAAATGTTTTTTAAGTAAATCAAGTAC 3’ and (positive strand) 5’CTCGATTCTATTAACAAGGG 3’, as described by Fey et al. (2013).

Fluorescently labeled strains were generated by transforming multicopy plasmids obtained from Dr. Marvin Whiteley’s lab (Georgia Institute of Technology), pCM29 (Pang et al., 2010) and pHC48 (Ibberson et al., 2016) into both JE2 and JE2 *gltT*::Tn via electroporation (Grosser & Richardson, 2016). This gave us the fluorescently labeled set of JE2.GFP, JE2.DsRed and JE2 *gltT*::Tn.GFP and JE2 *gltT*::Tn.DsRed (**Table 1**).

**Table 1.** List of *Staphylococcus aureus* strains used in this study. Plasmid pgltT was made as part of this study from the pOS1.plgT vector using methods described in Potter et al. (2020).

| Strain | Description | Plasmid | *gltT* Genotype | Reference |
| --- | --- | --- | --- | --- |
| JE2 | USA 300, background for NTML | None | wild type *gltT* | Fey et. al 2013 |
| PAO1 | *P. aeruginosa* strain |  |  |  |
| EV2 | Evolved JE2, *P. aeruginosa* tolerant isolate from EE1 | None | 1150 4bp deletion | This study |
| EV3 | Evolved JE2, *P. aeruginosa* tolerant isolate from EE1 | None | 951 G -> A early stop | This study |
| JE2 *gltT*::Tn | NTML transposon mutant NE566 transduced into JE2 background | None | *gltT*::Tn | Fey et. al 2013 |
| JE2 *gltT*::Tn (pgltT) | JE2 *gltT* transposon mutant complemented with pgltT construct | pgltT | complemented *gltT* | This study |
| EV2 (pgltT) | Evolved isolate EV2 complemented with pgltT construct | pgltT | complemented *gltT* | This study |
| JE2.GFP | JE2 fluorescently labeled with GFP | pCM29 | wild type *gltT* | Pang et al., 2010 |
| JE2.DsRed | JE2 fluorescently labeled with RFP | pHC48 | wild type *gltT* | Ibberson et al., 2016 |
| JE2 *gltT*::Tn.GFP | JE2 *gltT*::Tn fluorescently labeled with GFP | pCM29 | *gltT*::Tn | Pang et al., 2010 |
| JE2 *gltT*::Tn.DsRed | JE2 *gltT*::Tn fluorescently labeled with RFP | pHC48 | *gltT*::Tn | Ibberson et al., 2016 |

### Media

Cocultures for evolution experiments and phenotyping were conducted on TSA. After each coculture period each species was isolated on their respective isolation agar, Pseudomonas Isolation agar (PIA; BD Difco) and Staphylococcus isolation agar (SIA; BD Difco TSA with 7.5% NaCl). For liquid cultures, bacteria were cultured in lysogeny broth (LB; Teknova) which was supplemented with erythromycin (25 µg/mL) to select for transposon mutants and/or chloramphenicol (10 µg/mL) to maintain fluorescent plasmids. Chemically defined media with glucose (CDMG) was made according to Hussain et al. (1991), with varying levels of aspartate (1.1 mM or 2.2 mM) and glutamate (1.0 mM or 2.0 mM), as needed. CDMG batches were always used within 5 days and stored at room temperature, in the dark. Depleted Trypticase Soy Broth (TSB) medium, used for single-cell microscopy, was prepared by diluting an overnight culture of PAO1 1:100 into 10 mL TSB and growing the culture for either 3 or 16 hours before filter sterilizing (0.2 µm filter, Sarstedt) to remove cells from the supernatant.

### Experimental evolution

Before being cultured with *P. aeruginosa* PAO1, 4 single colony isolates of JE2 were picked and grown overnight in 3 mL of LB media in a test tube at 37°C in a rolling incubator. 10 µl of an overnight culture was then inoculated on 0.45 µm filters (MF-Millipore® Membrane Filter) on TSA and cultured at 37°C for 24 hours. Each filter was collected, and adhering cells were resuspended in 1.5 mL of LB media by vortexing for 30 seconds. To prepare *P. aeruginosa* for coculturing, a single colony of PAO1 was incubated in 3 mL of LB media overnight at 37°C in a rolling incubator. The optical density 600nm (OD_600_) of the resuspended *S. aureus* as well as the overnight PAO1 culture was measured. Cultures were normalized to the same OD by diluting the PAO1 overnight culture in LB before inoculating a coculture at a ratio of 30:1 (*S. aureus:P. aeruginosa).* A 30:1 inoculum ratio was determined through preliminary experiments to exert optimal amount selective pressure on *S. aureus* at without risking a population level extinction (approximately a 4-fold decrease in population size after coculture). 10 µl of coculture mixture was inoculated onto 0.45 µm filters on TSA plates and incubated at 37°C for 48 hours. To account for any adaptation to culture conditions, control populations of JE2 were passaged alongside experimental populations under identical conditions, but never cocultured with *P. aeruginosa*.

Each subsequent coculture was carried out by recovering filters and vortexing them in 1.5 mL of LB media for 30 seconds to collect adhering cells. Resuspended cell mixtures were serially diluted and plated for CFUs on SIA and PIA. 50 µl of resuspended coculture was also plated on SIA to be used to inoculate the next coculture (**Figure 1A**). After at least 24 hours of incubation at 37°C, isolated *S. aureus* was then collected off SIA plates with an inoculation loop and resuspended in LB media. This suspension was used to inoculate the next coculture period as well as to create a glycerol stock of the recovered population of *S. aureus*. PAO1 liquid cultures were incubated overnight at 37°C and then measured at OD_600_ and diluted to the same OD_600_ of the resuspended culture of *S. aureus* before being mixed at a ratio of 30:1. Inoculum densities fluctuated throughout the experiment based on the amount of *S. aureus* that could be recovered from the previous coculture period. Total inoculum densities ranged between 10^8^-10^10^ CFUs. We designed the experiment with an initial large population of *S. aureus* (10^8^-10^10^ CFUs) to mimic the event of *P. aeruginosa* invading an established *S. aureus* population as is often the case in CF-associated respiratory infections (Cystic Fibrosis Foundation, 2021).

**Figure 1.**
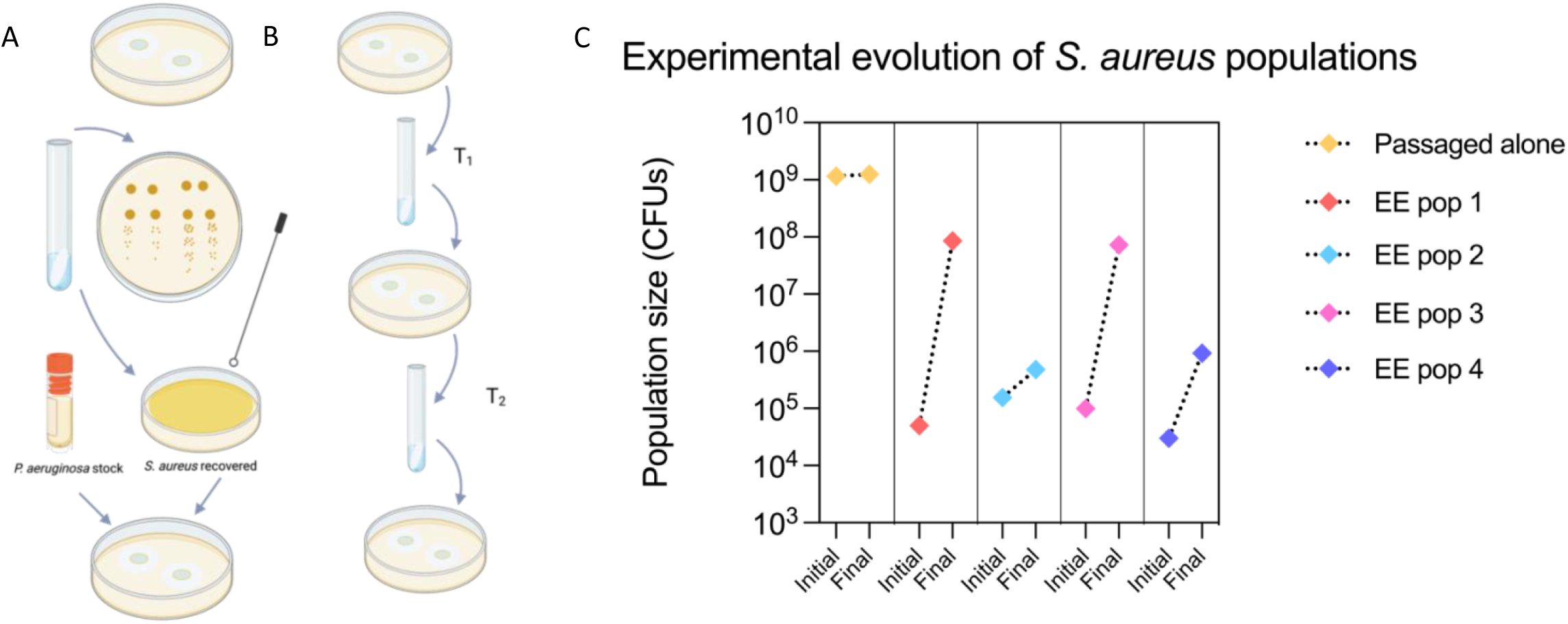
Experimental evolution with *Staphylococcus aureus* USA300 JE2 generates populations that are tolerant to *Pseudomonas aeruginosa* in coculture. *S. aureus* and *P. aeruginosa* were cocultured for 48 hours at an inoculation ratio of 30:1 (*S. aureus*:*P. aeruginosa*). A) Transfer procedure - cocultures are inoculated on solid TSA agar with 0.45 µm filter paper used to contain and recover the culture. After the coculture period, filters are resuspended in liquid media and serial dilutions of the resuspension are spot plated on selective agar. After 24 hours of growth, CFUs are counted for both species on their respective selective agar SIA and PIA. B) Serial transfer method - a new transfer occurs when *S. aureus* is isolated from the resuspension by plating 50 µl on SIA. After overnight incubation, *S. aureus* was inoculated with *P. aeruginosa* from a fresh overnight culture that was diluted to the same OD_600_. Control populations are repeatedly transferred under the same conditions but never exposed to *P. aeruginosa.* C) Experimental evolution results: four populations (pop 1-4) were evolved in the presence of PAO1 for 8 coculture periods; control population (passage alone) never exposed to *P. aeruginosa* also shown. CFU counts from the first coculture period (initial) and 8^th^ coculture period (final) are shown.

### Whole genome sequencing

Single colonies were isolated from evolved *S. aureus* populations and control populations and were used to inoculate overnight cultures and subsequent glycerol freezer stocks. Isolates were later struck out on SIA plates and incubated overnight at 37°C. Cells were collected off plates with an inoculation loop the next day and resuspended in 480 µl of EDTA. *S. aureus* cells were then lysed by adding 20 µl of 10 mg/mL lysozyme and 20 µl of 5 mg/mL lysostaphin to the resuspended cell mixture. This mixture was then incubated at 37°C for one hour before proceeding with the rest of the protocol outlined for the Promega Wizard genomic DNA purification kit (Silberstein et al., 2018). Genomic DNA was sequenced using the Illumina NextSeq 2000 platform at the Microbial Genome Sequencing Center (Pittsburgh, PA). Whole genome sequences were evaluated for quality using the program FASTQC (Wingett & Andrews, 2018) and adapter sequences were removed using Trimmomatic (Bolger et al., 2014). Sequences were then screened for variants using Snippy with JE2 NCBI NC_007793.1 sequence as a reference (Seeman, 2015).

### Complementation of *gltT*

The *gltT* gene was cloned using the multicopy pOS1 shuttle vector with the constitutive plgT promoter (Bubeck Wardenburg et al., 2006; Schneewind et al., 1992). The ancestral gene was amplified from wild-type JE2 using primers: 5’-AGAGCTCGAGATGGCTCTATTCAAGAG-3’ and 5’-AGATGGATCCTTAAATTGATTTTAAATATTCTTGAC-3’ and cloned downstream of the plgT promoter, as described in Potter et al. (2020). The resulting construct was confirmed by whole plasmid sequencing through Plasmidsaurus (Eugene, OR) and will be referred to as pgltT here (**Supplementary Figure 1**). The confirmed pgltT construct was transformed through electroporation into JE2 *gltT*::Tn, as well as the evolved isolate, EV2. All plasmids were transformed using electroporation (Grosser & Richardson, 2016)

### Analysis of variation of the *S. aureus gltT* gene

To detect the variation of the *gltT* gene, we assessed a dataset of *S. aureus* genomes that combined 380 assemblies from the Staphopia non-redundant diversity set (Petit III & Read, 2018) and 64 CF isolates (Bernardy et al., 2020) to create a dataset of 444 *S. aureus* genome assemblies. Then, we extracted and calculated the number of mutations in the genes *gltT, gltS, alsT, rpoD,* and *agrC,* using a custom software, LIVID (https://github.com/VishnuRaghuram94/LIVID), which performs *in-silico* PCR to extract nucleotide regions of interest from genome assemblies and compares the extracted sequence with a user-specified reference region to report mutations. For *gltT, gltS, alsT,* and *rpoD*, we used the corresponding gene sequences from the *S. aureus* strain JE2 (NCBI RefSeq accession GCF_002993865.1) as a reference. To account for different *agr* groups requiring a different reference sequence, we used the software tool AgrVATE (Raghuram et al., 2022) to calculate the number of mutations in *agrC*. Both LIVID and AgrVATE use Snippy v4.6 for identifying mutations (Seeman, 2015). LIVID was run with the parameters-x 1000 (minimum product size)-y 2000 (maximum product size)-d 5 (maximum allowed primer mismatch bases). AgrVATE was run with default parameters, as described in Raghuram et al. (2022) (**Supplementary Table 1**). Mutations labelled as ‘synonymous’ were single or multi-nucleotide substitutions that did not affect the amino acid sequence. Mutations labelled as ‘AA-sequence altering’ were single /multi-nucleotide substitutions in-frame insertions/deletions that cause local changes in the amino acid sequence. Putative ‘Loss of function’ variants include frameshift mutations, start-codon variants and early stops caused by non-synonymous mutations (**Supplementary Table 1**).

### Phenotypic testing for *P. aeruginosa* tolerance

*S. aureus* tolerance to *P. aeruginosa* strain PAO1, was determined by coculturing *S. aureus* and PAO1 at high initial densities (>10^8^ CFUs) at a 1:1 ratio for 24 hours and measuring recovered CFUs by serially diluting resuspended cultures and plating on SIA and PIA medias to select for *S. aureus* and *P. aeruginosa,* respectively (Bernardy et al., 2020). Phenotyping assays were carried out in 5 separate experiments with 2 biological replicates per strain.

### Murine acute pneumonia model

The impact of *gltT* activity on *S. aureus* colonization was determined in a murine acute pneumonia model. All animal procedures were conducted in accordance with the guidelines of the Emory University Institutional Animal Care and Use Committee (IACUC), under approved protocol number PROTO201700441. 8 to 10-week-old C57BL/6 female mice (Jackson Laboratories, Bar Harbor, ME) were anesthetized with a 0.2 mL mixture of ketamine (6.7 mg/mL) and xylazine (1.3 mg/mL) administered through intraperitoneal injection. All mice were euthanized 24 hours post-infection.

*S. aureus* strains JE2 and JE2 *gltT*::Tn were grown on SIA for 18 to 24 hours at 37°C and resuspended in phosphate buffered saline (PBS) to an OD_600_ of 8, corresponding to ∼2 x 10^9^ CFU/mL. Anesthetized mice were infected with 50 µl (∼1 x 10^8^ CFU) of *S. aureus* through intranasal administration (25 µL per nostril). Following euthanasia, whole lung and nasal wash were collected aseptically. The lungs were weighed and homogenized in 1 mL of PBS (Bullet Blender Storm 5). Homogenized lungs and nasal wash were serially diluted and plated on SIA to determine CFUs. For the acute pneumonia murine competition infection, JE2 and JE2 *gltT*::Tn were grown on SIA for 18 to 24 hours at 37°C and suspended in PBS to an OD_600_ of 14, followed by a 1:2 dilution corresponding to ∼4.8 x 10^9^ CFU/mL. Anesthetized mice were infected with 12.5 µl of culture of each *S. aureus* strain (∼6 x 10^7^ CFU) administered sequentially and single-strain control mice were infected with either 25 µl of JE2 or JE2 *gltT*::Tn (∼1 x 10^8^ CFU). Following euthanasia, whole lung and nasal wash were collected and processed following the procedures stated above. Serial dilutions were plated on both SIA and LA supplemented with erythromycin (25 µg/mL) to determine CFU, for both strains and JE2 *gltT*::Tn, respectively. Results were analyzed using one-way analysis of variance (ANOVA) corrected with Šidák in GraphPad Prism 9 (**Figure 4**).

### Competitive fitness assay

Fluorescently labeled JE2 and JE2 *gltT*::Tn were grown individually and together in complete CDMG (1.1 mM asp and 1.0 mM glu) (+A/+G) and CDMG with additional asp (2.2 mM asp) and no glu added (++A/0G). Cultures were inoculated at initial densities of OD_600_ 0.01 in flasks with 25 mL of media and incubated at 37°C with continuous shaking for 24 hours. CFUs were determined at inoculation, early growth (4 - 8 hours after inoculation) and endpoint (22 - 24 hours after inoculation). Both versions of each strain (JE2.GFP and JE2.DsRed and JE2 *gltT*::Tn.GFP and JE2 *gltT*::Tn.DsRed) were tested in these conditions in replicate experiments (**Figure 3**).

### Amino acid utilization

Amplite™ Fluorimetric L-Aspartate (Aspartic Acid) Assay Kit, and Amplite™ Fluorimetric Glutamic Acid Assay Kit *Red Fluorescence* (AAT Bioquest) were used to measure the concentration of aspartate and glutamate, respectively. Cell-free spent media was collected by filtering resuspended cocultures that were grown for 24 hours on TSA plates (as was done for phenotypic testing for *P. aeruginosa* tolerance) through 0.22 µm syringe filters. Spent media was collected from *S. aureus* monocultures and cocultures, as well as PAO1 monocultures and controls. Control conditions were made by inoculating sterile filters with 10 µl of LB or PBS and incubating for 24 hours. Measurements were taken across 3 separate experiments, each with 2 replicates for each culture condition.

### Single cell imaging

Batch cultures were grown in TSB media supplemented with chloramphenicol 10 µg/mL to maintain fluorescent plasmids. Overnight cultures were diluted 1:100 into 3 mL fresh TSB media and grown to mid-exponential phase in a test tube at 37°C in a shaking incubator (OD_600_ between 0.5-0.8). Subsequently, cells were washed 3 times with PBS to remove antibiotics and diluted to an OD_600_ of 0.1. Finally, a 1:1:1 coculture was prepared consisting of either JE2.DsRed + JE2 *gltT*::Tn.GFP + PAO1 or JE2.GFP + JE2 *gltT*::Tn.DsRed + PAO1. Agarose media was prepared by adding 1.5% agarose to either fresh or depleted TSB media.

A 1 mL droplet of agarose media was suspended between two coverslips and dried at room temperature for 30 minutes to create a ∼3 mm thick slap, which was then cut into 5×5 mm pads. 1 µL of the coculture was added to the pad and dried until the liquid was absorbed. Afterwards the pad was carefully inverted and placed in a glass-bottomed dish (WillCo Wells). 6 pads were added to the same dish with the following media conditions: fresh TSB, depleted TSB from a 3-hour culture, and depleted TSB from a 16-hour culture. Each media condition was inoculated with one of the two strain mixtures. A small piece of water-soaked tissue was added to preserve humidity and the dish was sealed with parafilm. The experiment was repeated four times using different biological replicates on two separate days. We conducted 4 replicate experiments with each labeled version (GFP or DsRed) of each strain being tested twice. Imaging of the samples began 1 hour after the agar pads were inoculated.

The pads were imaged using a Nikon Ti2 inverted microscope with perfect focus system, equipped with a Hamamatsu ORCA-Flash4.0 V3 Digital CMOS camera, a Nikon NA1.42 60X Plan Apochromat phase contrast oil objective, a SPECTRA-X LED fluorescent light source, and Chroma filter sets. Images were taken every 3 minutes in the phase, GFP, and RFP channels. Cells were kept at 37°C while imaging using a climate-controlled incubator (Oko-lab).

### Image analysis

To analyze colony growth, we segmented and tracked colonies using a custom-build pipeline (code is available at: https://github.com/simonvanvliet/PA-SA_Agarpads). The time-lapse movies were manually trimmed to remove later time points where cells overlapped in 3D or where excessive cell movement of *P. aeruginosa* was observed. Subsequently, images were registered using the phase_cross_correlation method of scikit-image (van der Walt et al., 2014). Segmentation of colony outlines was done using the Ilastik supervised pixel classification workflow (Berg et al., 2019). Pixels in the multi-channel image (phase, GFP, RFP) were classified in four classes (GFP / DsRed labeled *S. aureus*, *P. aeruginosa*, and background). The classification probabilities were post-processed using custom written python code to extract individual masks for each colony. In short, the probabilities were smoothed with a gaussian kernel and thresholded using a fixed threshold value of 0.5. The masks were then post-processed using a binary closing operation to fill-in gaps between neighboring cells. Finally, small objects were removed, and holes closed.

Colonies were tracked using a custom written tracking algorithm that matched colonies across subsequent frames based on the minimal center-to-center distance. Tracks were stopped when colonies merged. An automated filtering procedure was used to trim tracks whenever an unexpectedly large change in colony area was observed (indicative of missed colony merger and/or segmentation error).

Colony growth rate *r* was calculated as: 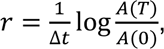, where *A*(0) and *A(T)* are the area (in pixels squared) of the colony at the start of the movie and after *T*=1 hour, and where Δ*t*=3 minutes is the imaging interval. To quantify the spatial arrangement, we calculated the minimal distance reached between the edge of the focal *S. aureus* colony to the closest pixel occupied by *P. aeruginosa*. Colonies within 300 pixels of the image frame were excluded from the analysis, as we could not accurately quantify their spatial arrangement.

## Results

### *gltT* truncation in *S. aureus* is an adaptation to *P. aeruginosa* tolerance

Serial transfer experimental evolution generated multiple populations of *S. aureus* with improved survival in the presence of *Pseudomonas aeruginosa* strain PAO1 (referred to as PAO1). At the completion of 8 serial transfers, two out of four experimental populations had significantly increased their relative survival compared to the JE2 ancestral strain. EE1 and EE3 both increased in the number of recovered CFUs by more than 3 orders of magnitude, with about 10^5^ CFUs being initially recovered to about 10^8^ CFUs recovered after 8 serial transfers (**Figure 1B**). Similar results were observed in replicate experiments. Single individual colonies from evolved *P. aeruginosa-*tolerant populations maintained this phenotype when cultured at a 1:1 ratio with PAO1 for 24 hours (**Supplementary Figure 2a**).

We sequenced the genomes of two single colonies from an evolved tolerant population EE1 and compared those sequences to those of colonies from an evolved but still sensitive population EE4 and colonies from control populations which were transferred in parallel with experimental populations but never cocultured with PAO1. Each isolate was screened for its survivability in coculture with PAO1 before being sequenced and the pattern of tolerance observed during the evolution experiment was confirmed. In the two colonies sequenced from population EE1, only one mutation site was unique to them, not appearing in any other compared sequence. Each evolved tolerant isolate (EV2 and EV3) had an independent putative loss of function mutation in the gene encoding for the *S. aureus* amino acid transporter, *gltT.* GltT has been previously described as being the sole aspartate transporter in *S. aureus* that also interacts with glutamate (Potter et al., 2020; Zeden et al., 2020; Zhao et al., 2018). In isolate EV3, a single nucleotide base substitution G → A introduced an early stop; in isolate EV2, a 4-base-pair deletion resulted in a frameshift mutation (**Figure 2A**). Both mutations occur between 800-1200bp downstream of the start codon and were predicted to truncate the protein by disrupting the 3’ portion of the coding region. Both single colony isolates displayed a *P. aeruginosa-*tolerant phenotype compared to their common ancestor, JE2 (**Supplementary Figure 2b**).

**Figure 2.**
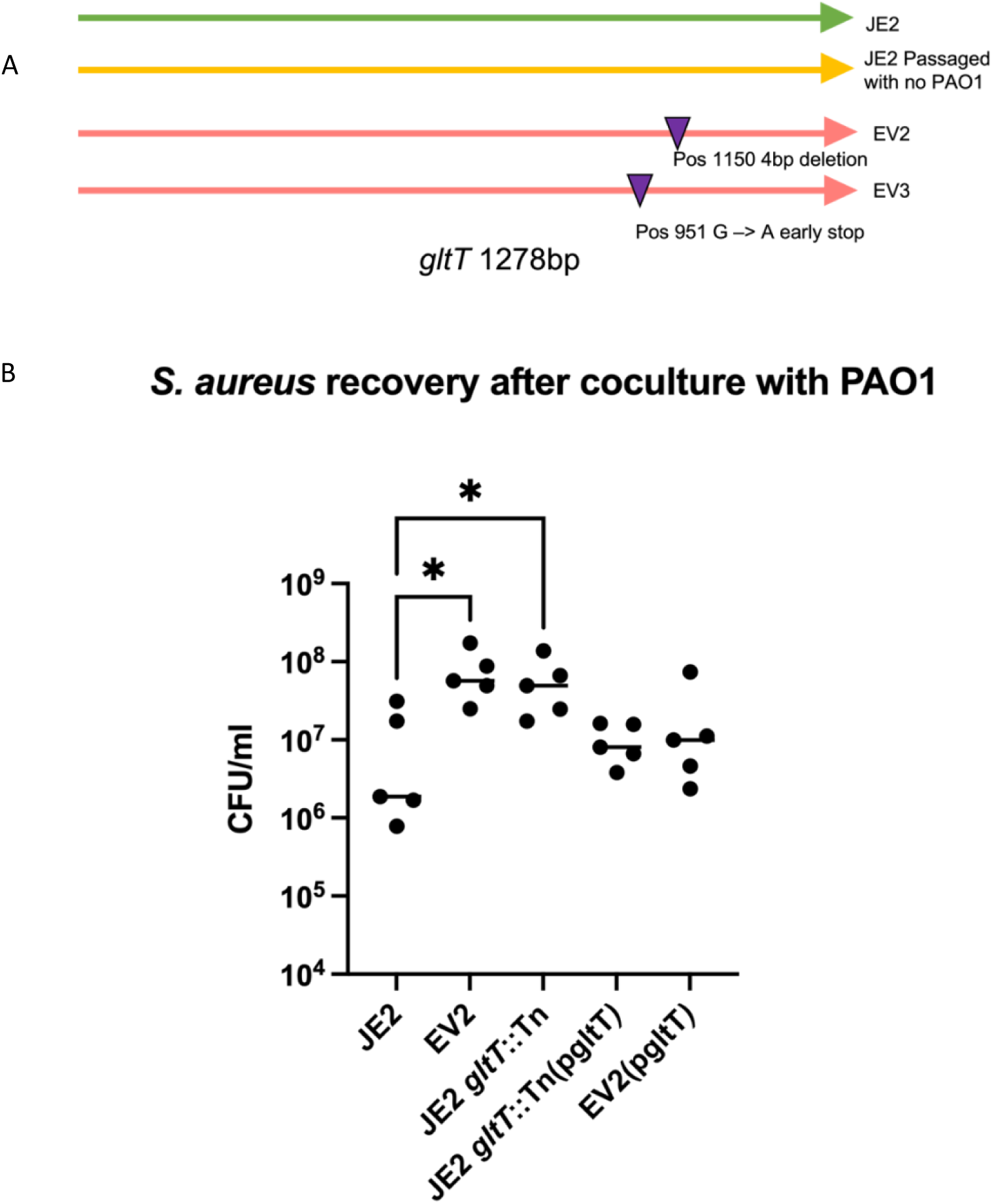
*gltT* truncation enhances *S. aureus* recovery after coculture with PAO1. A) Whole genome sequencing reveals two independent truncations of the aspartate transporter, *gltT*, in sequences of two single colony isolates EV2 and EV3 taken from the same evolved *P. aeruginosa-*tolerant population - EE pop 1 (Fig 1C). B) Evolved phenotype of *P. aeruginosa* tolerance is observed in the *gltT* transposon mutant JE2 *gltT*::Tn and evolved isolate EV2. Wild-type *P. aeruginosa* sensitivity is restored in the complemented transposon mutant JE2 *gltT*::Tn(pgltT) and complemented evolved isolate EV2(pgltT). Friedman’s test for multiple comparisons yielded p-values of 0.0127 and 0.0205 (*) when comparing JE2 CFUs to EV2 and JE2 *gltT*::Tn, respectively.

To confirm the linkage between *gltT* disruption and *P. aeruginosa-*tolerant phenotype, we retrieved the *gltT* mariner transposon knockout mutant, *SAUSA300_2329* from the Nebraska Transposon Mutant Library (NTML) (Fey et al., 2013). After transducing the mutation to the ancestral JE2 background, we validated the strain by PCR and confirmed that this strain, which we now refer to as JE2 *gltT*::Tn, had improved CFU recovery after coculture with PAO1 compared to its parent (**Figure 2B**). These results confirmed that the *gltT* disruption was responsible for the enhanced fitness of *S. aureus* in coculture with PAO1. When the mutant *gltT* strains were complemented in *trans*, the PAO1 sensitivity phenotype was restored for both the JE2 *gltT*::Tn and evolved isolate EV2 **(Figure 2B).**

### JE2 *gltT*::Tn outcompetes wild-type *S. aureus* in CDMG without glutamate

Growing strains individually in CDMG (**Supplementary Table 2**) or rich LB (**Supplementary Figure 3**) media yielded no insights into fitness differences associated with *gltT* disruption. However, we hypothesized that *gltT* mutants may be able to outcompete wild-type *S. aureus* under certain conditions. To test this, we conducted competition experiments with mutant and wild type strains fluorescently labeled with either GFP or DsRed multicopy plasmids. Labeled strains were inoculated in complete CDMG with 1.1 mM asp and 1.0 mM glu (+A/+G) and CDMG with double the amount of aspartate (2.2 mM asp) and no glutamate added (++A/0G). CFU counts from three replicate experiments showed that JE2 and JE2 *gltT*::Tn were equally fit when grown in complete CDMG, with each strain making up about half of the total culture density. Total CFUs were greater than 10^8^ CFUs across all conditions and replicates. However, in the (++A/0G) condition, JE2 *gltT*::Tn outcompeted wild-type JE2 which only made up about 5% of all CFUs recovered from this growth condition (**Figure 3**).

**Figure 3.**
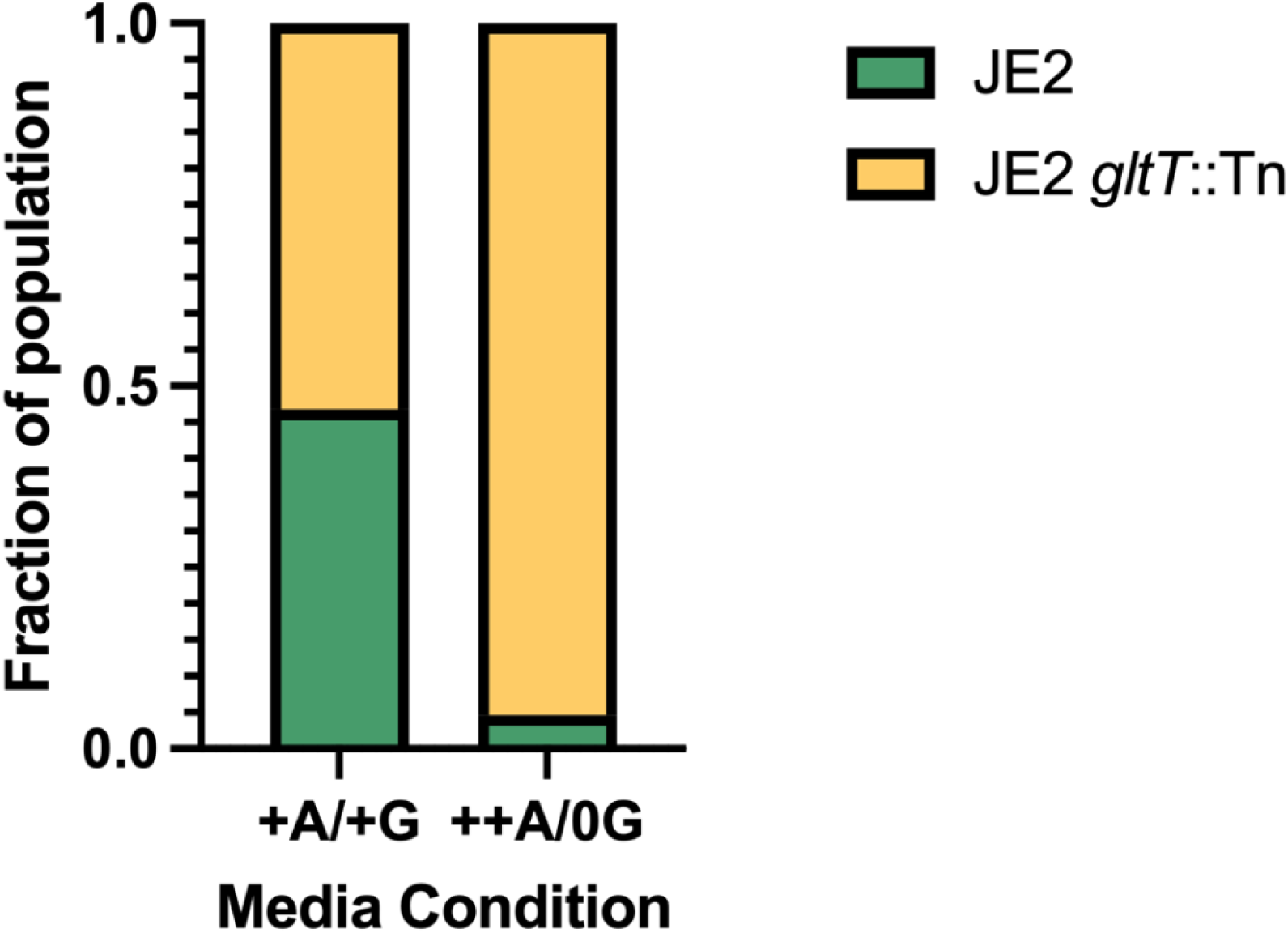
JE2 *gltT*::Tn outcompetes wild-type JE2 in CDMG when glutamate is limiting and aspartate is in excess. JE2.GFP, JE2.DsRed and JE2 *gltT*::Tn.GFP and JE2 *gltT*::Tn.DsRed strains labeled with fluorescent plasmids were used to test for competitive fitness in CDMG conditions. One GFP labeled strain and one ds.Red labeled strain were cultured for 24 hours in CDMG media alone and in coculture. Incubated cultures were then serially diluted and incubated overnight at 37°C before CFUs were counted and number of red and green colonies recorded. Each version of each strain was tested at least once across 3 biological replicates. JE2 *gltT*::Tn made up the majority of CFUs recovered after 24 hours when aspartate (A) was in excess (++), and glutamate (G) was limited (0G) (++A/0G) Chi-square p-value< 0.0001.

### Growth rate differences, as measured by single cell microscopy, are not responsible for the *P. aeruginosa*-tolerant phenotype

Based on results from CDMG assays, we hypothesized that glutamate depletion by PAO1 would reduce the growth rate in nearby JE2 cells, but not in JE2 *gltT*::Tn cells. To test this hypothesis, we conducted single cell microscopy with wild-type JE2 and JE2 *gltT*::Tn and measured fitness by CFU counts after a 24 hour coculture with PAO1. We conducted single cell image analysis on cocultures with equal starting ratios of JE2, JE2 *gltT*::Tn and PAO1 using agar pads and GFP and DsRed fluorescently labeled strains. We hypothesized that growth rate differences may only be apparent in depleted media conditions as CFU differences were most obvious between JE2 and JE2 *gltT*::Tn when cocultures are inoculated at high initial densities. However, even in depleted TSB collected from a 16 hour culture of PAO1, JE2 *gltT*::Tn did not have an observable difference in growth rate compared to wild-type JE2 (**Figure 4**). Moreover, we did not find a dependence of *S. aureus* growth rates for either wild-type JE2 or JE2 *gltT*::Tn based on their proximity to PAO1 colonies (**Supplementary Figure 4**). In fact, the only growth difference observed was a slight advantage for wild-type JE2 in depleted TSB collected from a 3-hour culture of PAO1. Both strains had very low growth rates in the most depleted media condition, TSB collected from a 16-hour culture of PAO1. Morphology of microcolonies were indistinguishable between *S. aureus* strains and no evidence of small colony variants were observed. These results suggest that the growth advantage of JE2 *gltT:*:Tn is a population-level trait that is not explained by growth rate differences between it and its isogenic wild-type counterpart.

**Figure 4.**
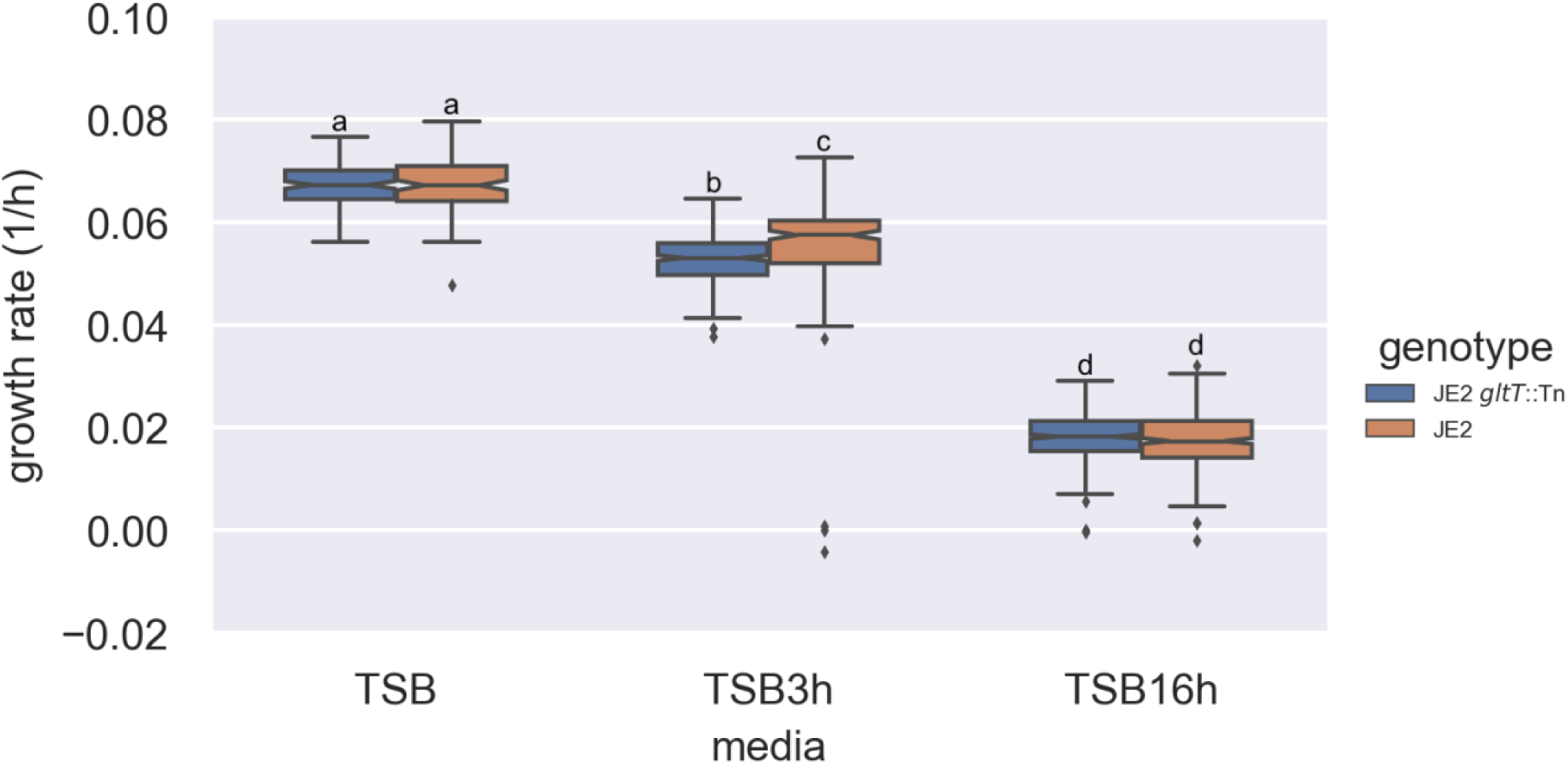
Growth rates of wild-type JE2 and JE2 *gltT*::Tn in coculture with PAO1 in rich and depleted media. Fluorescently labeled strains - JE2.GFP or JE2.DsRed and JE2 *gltT*::Tn.GFP or JE2 *gltT*::Tn.DsRed were cultured together with PAO1 on agar pads made from rich Trypticase Soy Broth (TSB) and depleted cell-free TSB collected from 3 hour and 16-hour PAO1 monocultures. Growth rates of micro colonies were measured using single cell microscopy image analysis. Wild-type JE2 is observed to have a slight growth advantage over mutant JE2 *gltT*::Tn in slightly depleted TSB from a 3-hour culture, however strains grow generally at the same rate in all other conditions. Data was collected over four replicates for each of the two strain combinations. We did not observe consistent differences between the different DsRed or GFP strain combinations and their data was thus pooled. PAO1 also carried a GFP label and was easily distinguished based on cell shape. The two-way ANOVA revealed significant effects of media (F(2, 18) = 11800.29, p < 0.001) and genotype (F(1, 18) = 4.37, p = 0.036), as well as a significant interaction between media and genotype (F(2, 18) = 17.55, p < 0.001) on growth. Post-hoc Tukey’s HSD test indicated significant differences between different media conditions (p < 0.05), suggesting that the growth varied significantly depending on the media used. Additionally, significant differences were observed between genotypes in the TSB3h media (p < 0.05).

### *gltT* disruption alters amino acid uptake in *S. aureus* strains

We measured concentrations of aspartate and glutamate in cell-free spent media that was collected from 24-hour TSA cultures (monocultures and cocultures with PAO1) of the following *S. aureus* strains: JE2, JE2 *gltT*::Tn, EV2 and complemented strains JE2 *gltT*::Tn (pgltT) and EV2 (pgltT). These data revealed that aspartate concentration remained highest in all culture conditions where PAO1 was not present (**Figure 5**). In the case of glutamate, there was very little of the amino acid remaining in any culture condition with wild-type or *gltT* complemented *S. aureus* strains even when PAO1 was not present. Both amino acids were present at higher concentrations in the spent media of *S. aureus* monocultures compared that of cocultures or PAO1 monocultures. This suggested that PAO1 was more efficient at metabolizing both aspartate and glutamate than *S. aureus.* Additionally, the levels of remaining glutamate were higher in *S. aureus* cultures of the *gltT* mutant, suggesting that a functional *gltT* locus was essential to utilizing most of the available glutamate (**Figure 5**).

**Figure 5.**
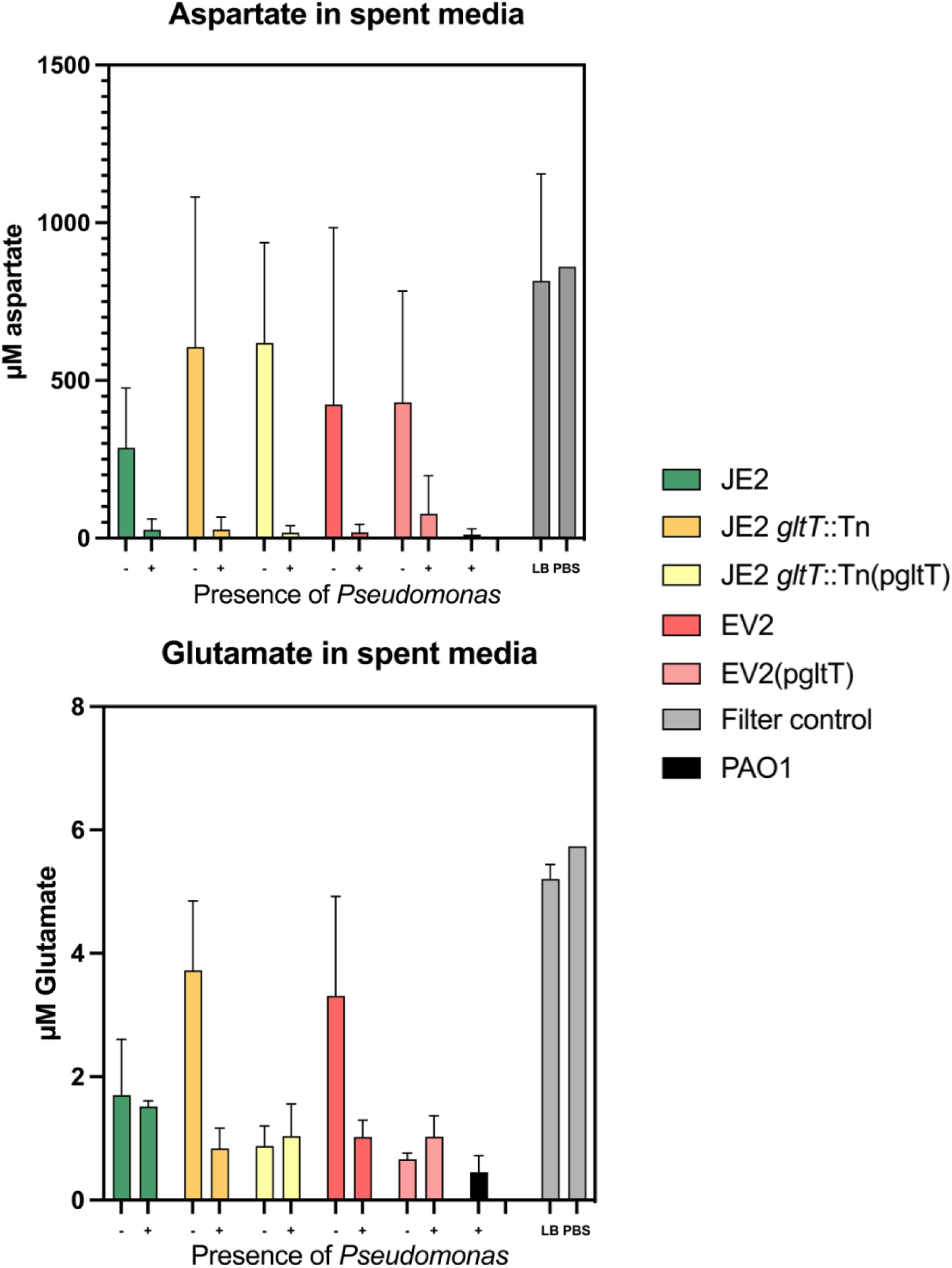
*gltT* is required for *S. aureus* glutamate uptake and aspartate and glutamate are greatly reduced when PAO1 is present. Remaining aspartate and glutamate levels after 24-hours of culturing on 0.45µM filters on TSA plates, were measured using Amplite Fluorimetric kits. Both cocultures with PAO1 (+) and *S. aureus* monocultures (-) were tested. Control conditions were tested by inoculating filters with 10µL of LB media or 10µL of PBS.

### *gltT* disruption does not impact *S. aureus* host colonization

To gain an understanding of the impact of *gltT* disruption in a host environment, we carried out experiments using an acute murine pneumonia model system where mice were infected with JE2 *gltT*::Tn, wild type JE2, or both strains in a coinfection. There were similar population sizes of the ancestral JE2 strain and JE2 *gltT*::Tn recovered from both the nasal wash and lung tissue for both single strain cultures and cocultures of the two strains **(Figure 6).** Therefore, we concluded that *gltT* disruption did not greatly impact *S. aureus’* ability to colonize host respiratory tissues.

**Figure 6.**
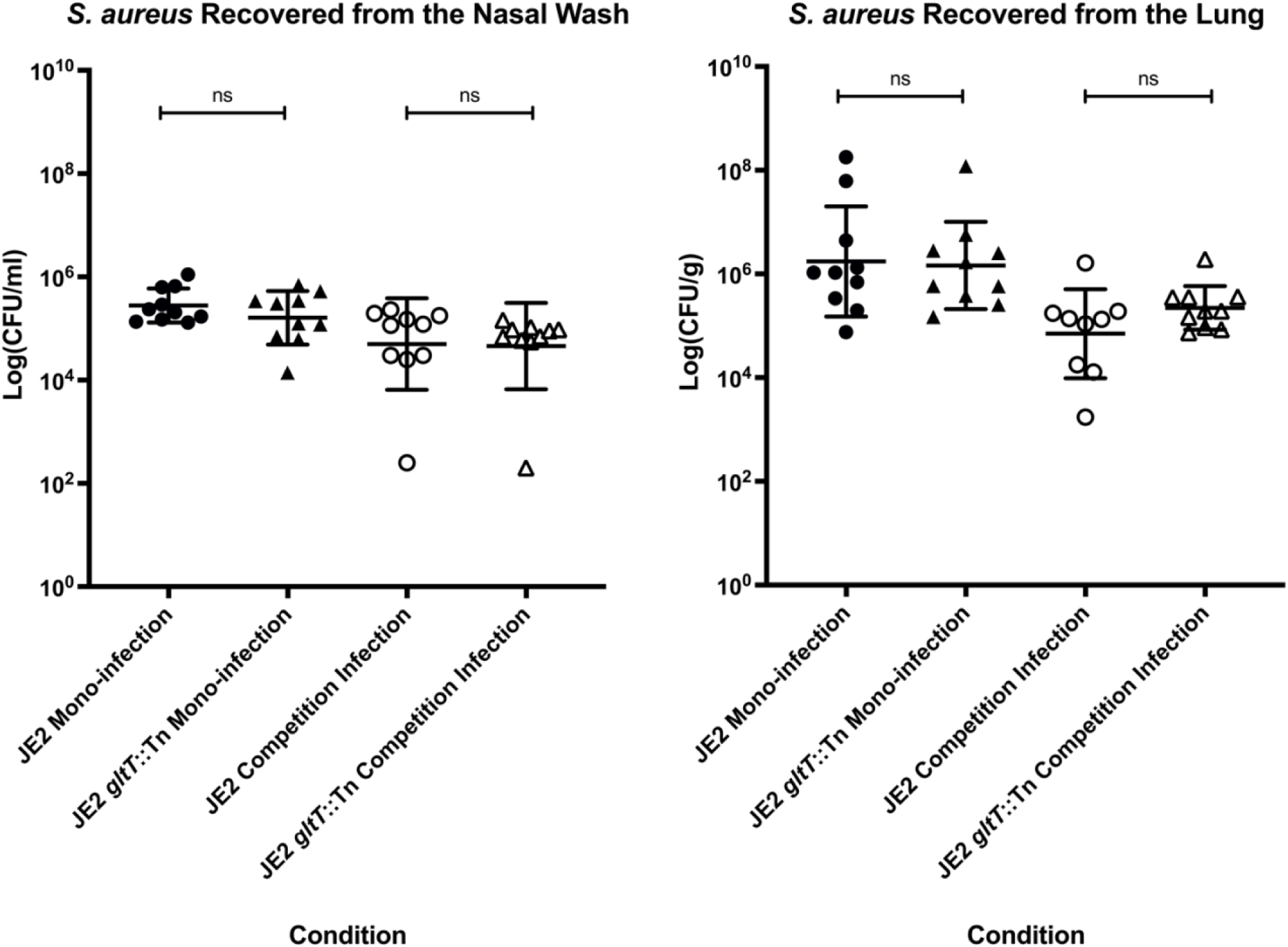
Colonization ability of *S. aureus* is not impacted by *gltT* genotype. Similar amounts of *S. aureus* are recovered from both lung (left) and nasal wash (right) when JE2 or JE2 *gltT*::Tn are cultured in a mouse alone or in coculture. All mice were euthanized 24 hours post infection. All statistics were performed using GraphPad Prism 9 using one-way analysis of variance (ANOVA) with Šidák correction. Similar levels of colonization in the lungs and upper respiratory tract (nasopharynx) was observed between JE2 and JE2 *gltT*::Tn in 8-to 10-week-old C57BL/6 female mice. For mono-infections, 1×10^8^ CFU of JE2 and JE2 *gltT*::Tn was administered intranasally. For the competition infections, 6×10^7^ CFU of both JE2 and JE2 *gltT*::Tn was sequentially administered intranasally. After 24-hours post-infection, all mice were euthanized and CFUs from the nasal wash and lungs were recovered either on SIA (mono-infection) or both LB + erythromycin (25 µg/ml) and SIA (competition infection). All statistics were performed using GraphPad Prism 9 using one-way analysis of variance (ANOVA) with Šidák correction.

### *gltT* disruption was rare in diverse *S. aureus* genomes

Findings from the evolution experiment and *gltT* mutant phenotyping tests indicated that *gltT* could be disrupted without severe impacts on strain fitness. We therefore, sought to estimate the variability of *gltT* across diverse *S. aureus* lineages, including CF-associated isolates. Previous work had shown that *gltT* is a core gene (Petit III & Read, 2018) and so we were able to look at *gltT* variability by screening a diverse dataset of 444 *S. aureus* genomes representing 380 MLST sequence types. We did not find mutations identical to the ones we saw during our experimental evolution. Furthermore, we identified only one mutation that caused an early stop, at position 1255, truncating the protein by 8 amino acids. This was the only putative non-functional mutant we observed in *gltT* and it was present in only one sample. This putative loss of function mutant was not isolated from a CF-associated infection.

We did not observe a significant enrichment of mutations in *gltT* when compared to other amino acid transporters in our dataset. We observed 115 occurrences of non-synonymous mutations in *gltT*, with 26 distinct mutations, 25 of which were found in < 10 strains. One mutation - a glutamate → aspartate change at position 891, was found in in 71 strains. Overall, these results suggested that *gltT* disruption was rare in *S. aureus*, even compared to other core genes encoding amino acid transporters (**Supplementary Table 1**). This was also true for CF associated isolates which, in our screen, did not have elevated rates of mutation in *gltT* compared to non-CF isolates.

## Discussion

### Impact of inactivation of *S. aureus gltT* gene in *S. aureus-P. aeruginosa* interactions

Interactions between *S. aureus* and *P. aeruginosa* have proven to be complex, and dependent on environment and strain background (Bernardy et al., 2020, 2022). Studies have implicated factors such as the *P. aeruginosa* mucoidy phenotype, *Pseudomonas* excreted compounds or toxins, and *S. aureus* metabolic pathways such as the production of acetoin, as important factors in the interspecific interactions between *S. aureus* and *P. aeruginosa* (Barraza & Whiteley, 2021; Bernardy et al., 2020, 2022; Camus et al., 2020; Lasse et al., 2022; Zarrella & Khare, 2021). Surprisingly, despite the wealth of research on the coexistence and competition of these species, in our experimental system, we observed mutation of a highly conserved gene, *gltT*, that had not previously been linked to *S. aureus-P. aeruginosa* co-occurrence. Our findings indicated that *S. aureus* JE2 and *P. aeruginosa* PAO1 directly compete over limiting glutamate, particularly when grown at high densities on TSA plates. Our evolved isolates appear to have gained a phenotypic advantage over their JE2 ancestor by disrupting the *gltT* locus – limiting import of aspartate and glutamate and likely relying on alternative metabolic pathways. Under the conditions of the evolution experiment we designed here, there is apparently significant selective pressure for *S. aureus* to optimize its amino acid metabolism for a glutamate-limited environment. The *S. aureus gltT* gene was also found to be disrupted in osteomyelitis studies that revealed how excess glutamate competitively inhibits aspartate transport through *gltT* (Potter et al., 2020). While the osteomyelitis study of Potter et al. (2020)presents the inverse of the nutrient landscape *S. aureus* is adapting to in our experiment – a challenge of excess glutamate rather than it being a limiting nutrient - it also demonstrates the importance of exogenous amino acids in *S. aureus* competitive fitness and the importance of altering metabolic pathways as an adaptive strategy in changing environments. Our findings suggest that we still do not understand enough about the interaction between *S. aureus* and *P. aeruginosa* to predict the genes that give adaptive advantages in any given combination of strains and environmental conditions.

We acknowledge some limitations of our experimental evolution approach. For instance, while fresh PAO1 was introduced to each coculture period we found that a minority population of *P. aeruginosa* were able to survive on SIA agar even though they did not form colonies. We suspect that some *P. aeruginosa* cells may have been transferred between coculture periods and could have coevolved with the JE2 populations. However, fresh PAO1 was introduced at each coculture period, and the evolved *P. aeruginosa-*tolerant phenotypes of *S. aureus* were maintained after populations were fully isolated from any retained PAO1. Therefore, any effect of carryover *P. aeruginosa* appeared to have little effect compared to the larger population of fresh introduced ancestral PAO1. Additionally, while the population sizes in our evolution experiment are likely much denser than what occurs during a CF lung infection, populations of *S. aureus* and *P. aeruginosa* have both been observed at levels as high as 10^8^ CFUs/mL in CF sputum samples collected from coinfected patients (Fischer et al., 2020)

### The role of aspartate and glutamate in *P. aeruginosa* tolerance

Previous analysis of *S. aureus* metabolism has shown that glutamate derivatives are required for *S. aureus* to metabolize aspartate into oxaloacetate, a secondary metabolite required in the citric acid cycle (Halsey et al., 2017). Therefore, the absence of extracellular glutamate may lead to reduced activity of the TCA cycle. This could explain why all tested *S. aureus* strains showed a reduced growth-rate when glutamate was not present in CDMG compared to complete CDMG (**Supplementary Table 2**). However, we did not observe significant growth rate differences between wild type JE2 *S. aureus* and JE2 *gltT*::Tn in our CDMG monoculture assays or by single-cell microscopy. These data suggest that despite its important role in amino acid metabolism there were few apparent fitness trade-offs associated with disrupting *gltT* when *P. aeruginosa* is not present in standard laboratory conditions (Zhao et al., 2018). In addition, the finding that *gltT* mutations are extremely rare in non-laboratory adapted stains reinforces the key metabolic role of this core gene. We postulate that continuing to import aspartate in the wild-type strain in the absence of glutamate may lead to a buildup of aspartate intracellularly and a corresponding reduced competitive fitness. This hypothesis is supported by our finding that along with its increased *P. aeruginosa*-tolerance, JE2 *gltT*::Tn was able to outcompete wildtype JE2 in the CDMG condition where additional aspartate was added and no glutamate was added (++A/0G) **(Figure 3)**.

Further experimentation is needed to better understand how amino acid metabolism facilitates *S. aureus-P. aeruginosa* interactions; however, we conclude here that in our evolution experiment, *S. aureus* is primarily adapting to the limitation of glutamate in its environment by disrupting its aspartate transporter and relying on alternative metabolic pathways to carry out the TCA cycle. In short, the major source of negative selective pressure that *S. aureus* experienced when grown in the presence of *P. aeruginosa* was competition over exogenous amino acids.

### Experimental evolution as a useful tool for studying pathogens

Experimental or directed evolution experiments carried out in laboratory conditions can be a powerful way to reduce complex adaptive phenotypes in important pathogens like *S. aureus* and *P. aeruginosa* to single genetic determinants. The lab environments used for experimental evolution studies are removed from the conditions we study, such as the cystic fibrosis lung environment or the host environment in general. Despite this however, there is still a lot to be gained from evolution experiments conducted in laboratory conditions with lab-adapted strains. Even if such experiments identify genes that are highly conserved (and thus unlikely to be important for adaptation in the setting of human infection) as we have found in this study, these findings reveal potential adaptive trajectories that may lead to possible treatment targets as well as a greater understanding of pathogen biology and physiology. For instance, our findings here lay important groundwork in the development of coinfection disruption therapy by highlighting the importance of the nutrient landscape in the facilitation of *S. aureus-P. aeruginosa* coexistence in the cystic fibrosis lung or other chronic infections. The link between *gltT* and *P. aeruginosa* tolerance likely could not have been identified by screening clinical isolates because the gene is so highly conserved. Our *in vitro* experiments suggest that *gltT* mutants can colonize lung tissue just as well as wild-type strains and would be more likely to coexist with *Pseudomonas* in a coinfection. Therefore, even though likely rare, this genotype could be important to screen for when treating coinfections and is certainly important to consider in the development of new therapies to treat polymicrobial infections.

## Acknowledgements

This work was supported in part with grants for the National Institutes of Health (R21 AI148847) and the US Cystic Fibrosis Foundation (GOLDBE19I0) to JBG. Research reported in this publication was supported in part by National Institute of Allergy and Infectious Diseases of the National Institutes of Health (T32AI138952), and Emory University and the Infectious Disease Across Scales Training Program (IDASTP) (AMA). The content is solely the responsibility of the authors and does not necessarily represent the official views of the National Institutes of Health, Emory University or IDASTP.

GB and SVV were supported by an Ambizione grant from the Swiss National Science Foundation (grant no. PZ00P3_202186) and by the University of Basel.

We would like to thank Rachel Done for her assistance with cloning and strain preparation. We would also like to thank members of the Goldberg and Read labs for their support and review of the manuscript.

